# Probabilistic risk assessment of pesticides under future agricultural and climate scenarios using a Bayesian network

**DOI:** 10.1101/2022.05.30.493954

**Authors:** Sophie Mentzel, Merete Grung, Roger Holten, Knut Erik Tollefsen, Marianne Stenrød, S. Jannicke Moe

**Affiliations:** Norwegian Institute for Water Research, Section for Ecotoxicology and Risk Assessment, Oslo, Norway; Norwegian Institute of Bioeconomy Research, Division for biotechnology and plant health, Ås, Norway; Norwegian University of Life Sciences (NMBU), Ås, Norway

## Abstract

The use of Bayesian networks (BN) for environmental risk assessment has increased in recent years. One reason is that they offer a more transparent way to characterize risk and evaluate uncertainty than the traditional risk assessment paradigms. In this study, we explore a new approach to probabilistic risk assessment by developing and applying a BN as a meta-model for a Norwegian agricultural site. The model uses predictions from a process-based pesticide exposure model (World Integrated System for Pesticide Exposure - WISPE) in the exposure characterization and species sensitivity data from toxicity tests in the effect characterization. The probability distributions for exposure and effect are then combined into a risk characterization (i.e. the probability distribution of a risk quotient), which is a common measure of the exceedance of an environmentally safe exposure threshold. In this way, we aim to use the BN model to better account for variabilities of both pesticide exposure and effects to the aquatic environment than traditional risk assessment. Furthermore, the BN is able to link different types of future scenarios to the exposure assessment, taking into account both effects of climate change on pesticides fate and transport, and changes in pesticide application. We used climate projections from IPCC scenario A1B and two global circulation models (ECHAM5-r3 and HADCM3-Q0), which projected daily values of temperature and precipitation for Northern Europe until 2100. In Northern Europe, increased temperature and precipitation is expected to cause an increase in weed infestations, plant disease and insect pests, which in turn can result in altered agricultural practices, such as the use of new crop types and changes in pesticide application patterns. We used the WISPE model to link climate and pesticide application scenarios, environmental factors such as soil properties and field slope together with chemical properties (e.g. half-life in soil, water solubility, soil adsorption), to predict the pesticide exposure in streams adjacent to the agricultural fields. The model was parameterized and evaluated for five selected pesticides: the herbicides clopyralid, fluroxypyr-meptyl, and 2-(4-chloro-2-methylphenoxy) acetic acid (MCPA), and the fungicides prothiocanzole and trifloxystrobin. This approach enabled the estimation and visualization of probability distribution of the risk quotients representing the alternative climate models and application scenarios for the future time horizons 2050 and 2075. The currently used climate projections resulted in only minor changes in future risk directly through the meteorological variables. A stronger increase in risk was predicted for the scenarios with increased pesticide application, which in turn can represent an adaptation to a future climate with higher pest pressures. Further advancement of BN modelling as demonstrated herein is anticipated to aid targeted management of ecological risks in support of future research, industry and regulatory needs.

## 1 Introduction

Climate change (CC) is expected to shift weather patterns, and in sequence can alter the way water and food resources are obtained and managed worldwide by 2050. Already today, European assessment for rivers and lakes report that 5 – 15 % of the monitoring stations show exceedances of environmental quality standards by herbicides, and 3 – 8 % by insecticides over the period 2007-2017 (Mohaupt et al., 2020). Albeit, in future pesticides will be extensively used as they will continue to play a vital role in the food production process and food security (Popp et al., 2013). Despite thorough regulation of pesticides, large knowledge gaps continue to hinder risk assessment, especially when it comes to potential environmental impact of pesticide mixtures and impacts of climate and regional factors (Weisner et al., 2021, Topping et al., 2020). In Northern Europe, an predicted increase in plant diseases and insect pests may consequently lead to higher pesticide use and thereby occurring concentration of pesticides in the environment (Kattwinkel et al., 2011, Delcour et al., 2015, Sutherst et al., 2011). As pesticide environmental fate and exposure scenarios deviate from EU predictions due to spatial (regional) or temporal differences (Holten et al., 2018, Stenrød et al., 2008), for Norway and the Nordic countries, the pesticide use, emissions, exposure and fate are not adequately represented by the standardized EU model scenarios (Stenrød et al., 2016). To safeguard environment health better, there is a need to improve the integration of trend connected to CC into environmental risk assessments of pesticides, such as the shifts in climate conditions or application patterns. This should subsequently enable better informed risk management and thus minimize harmful effects to the environment by enhancing decision making for mitigation efforts.

Current paradigms for environmental risk assessment (ERA) of pesticides typically aim to take into account the variability of species sensitivities by estimating a proportion of affected species in a community, which is used to define a predicted no-effect concentration (PNEC) of the pesticide (More et al., 2019). The traditional risk characterization of pesticides usually uses single-value e.g. Toxic exposure ratio derived from predicted environmental concentration (PEC) divided by PNEC to assess whether a chemical substance poses a risk to the environment (EC, 2011). In this study we applied a more general approach using a risk quotient (RQ) that is calculated as PEC/PNEC, where a potential risk to the environment is assumed whenever the safe concentration (PNEC) exceeds the PEC (Bruijn et al., 2002, More et al., 2019). These derived point estimates may convey an unjustified sense of accuracy (Rai et al., 2002), as they do not consider the variability of pesticides concentrations in the environment or other factors that influence the exposure of biota to these chemicals. Especially in Europe, these traditional methods seek to avoid underestimating risk by using conservative assumptions (i.e. assessment factors) to account for various sources of uncertainty (Verdonck, 2003). This way, protective decision making relies on precautionary safety margins (Fairbrother et al., 2015). Spatial and temporal variations in exposure are caused by many factors, including changing environmental characteristics and contamination sources (Artigas et al., 2012) that can cause uncertainty in applied scenarios. There is a need for risk assessment to better account for uncertainty and variability in chemical exposure (Belanger and Carr, 2020).

Probabilistic risk assessment make use of probability distributions to characterize uncertainty in all parts of the risk characterization (Mentzel et al., 2021, EUFRAM, 2006). Ergo, fully probabilistic risk characterization can better account for spatial and temporal variability of both chemical concentrations and species sensitivity (EUFRAM, 2006, Verdonck, 2003, Fairbrother et al., 2015, Solomon et al., 2000). Several probabilistic methods have been proposed to characterize risk while including estimation of stochastic properties and uncertainty (Solomon et al., 2000). The general responsibility of scientists to communicate uncertainties has also been highlighted by the EU (EFSA and BfR, 2019). Already two decades ago, the use of probabilistic risk assessment has been recommended for the European Union (EU) (Jager et al., 2001) but is still not commonly applied in regulatory risk assessment (Fairbrother et al., 2015). Probabilistic methods that incorporate distributions for exposure and effect are e.g. joint probability curves and quantitative overlap. Generally, probabilistic methods require more data for calculation of distributions compared to traditional ERA, but on the other hand probabilistic methods make better use of available data (Campbell et al., 2000, Verdonck, 2003), but generally make better use of available data. These methods consider multiple possibilities and are able to produce outputs with more ecological meaning (FOCUS, 2007). Nevertheless, some of the results are difficult to communicate and thereby challenging for decision-makers to interpret and understand (Verdonck, 2003, FOCUS, 2007), possibly because they are often based on cumulative distribution curves (EUFRAM, 2006). We propose the use of Bayesian networks (BN) as a more user-friendly and intuitive method for probabilistic risk assessment of pesticides (Mentzel et al. 2021). To demonstrate this, we developed a BN model to assess environmental risk of pesticides under future scenarios. Therefore, we explored an approach that’s includes the output of a pesticide exposure prediction platform for a representative Northern European area (WISPE; Bolli et al. (2013)) under different climate and pesticide use scenarios. The first aim of this study was to incorporate these various scenarios into a probabilistic approach of risk characterization and secondly to link the predicted environmental concentrations from the model with toxicity data. From this we present the results from a risk characterization for a Norwegian case study including an evaluation of direct and indirect effects of CC.

## 2 Material and Method

### 2.1 Approach

#### 2.1.1 Bayesian network model, structure and implementation

Bayesian methods have been recommended for uncertainty analysis in the process of identifying limitations in scientific knowledge and evaluating their implications for scientific conclusion by the European Food Safety Authority (EFSA et al., 2018). Bayesian networks (BNs) are a branch of Bayesian approaches that have been increasingly used in environmental risk assessment and management (Moe et al., 2021b, Aguilera et al., 2011, Kaikkonen et al., 2021). BNs are probabilistic and graphical models, more specifically directed acyclic graphs (DAG) (Kanes et al., 2017) that have no feedback loops (Troldborg et al., 2021). The nodes (variables) are connected through links (potentially causal relationships) shown as arcs representing conditional probability tables (CPTs) (Kjærulff and Madsen, 2013). Each node has states (typically intervals) that are quantified by probability distributions. To update the probability distributions of network, the Bayes’ rule is implemented to combine prior probabilities with new evidence (Carriger et al., 2016). One of the main benefits of BNs is that all components can be quantified by probability distributions, and probabilistic risk calculation can be carried out. Along these lines, BNs can incorporate various sources of information such as expert opinion, literature and model outputs, enabling a greater use of available data and knowledge (Carriger et al., 2016, Carriger and Newman, 2012). A recent example of implementation of BNs in risk assessment by Carriger and Barron (2020) showed the calculation of a probabilistic risk quotient (RQ) by combining probability distributions of exposure and effects of mercury for the Florida panther. Another example for the application of spatial BNs for probabilistic exposure assessment of pesticide on a field level has been carried out by Troldborg et al. (2021). One more example focused on the risk estimation to the aquatic environment by pesticides using the BN to quantify the RQ and associated uncertainties by Mentzel et al. (2021). This approach used exposure and effect distributions to derive a RQ distribution.

The BN conceptual model developed here is based on Mentzel et al. (2021) and consist of four modules: (1) future scenarios (orange), (2) pesticide exposure (blue), (3) toxic effect (green) and (4) risk characterization (grey) (Figure 1). The scenario module contains a scenario node that is based on the climate and pesticide application. These scenarios determine the instantaneous pesticide concentration and its probability distribution (pesticide exposure module). This instantaneous node together with the set time since application (node) determines the distribution of the time-specific concentration node, and a linear equation was used in the calculation. The risk characterization module composes the exposure/effect ratio node that together with an appropriate precautionary factor predicts the probabilities of the RQ intervals. The finalized BN can be instantiated by selecting a scenario and specifying the time since application of interest as evidence. Given this evidence, probability distributions will be updated throughout the network.

**Figure 1.**
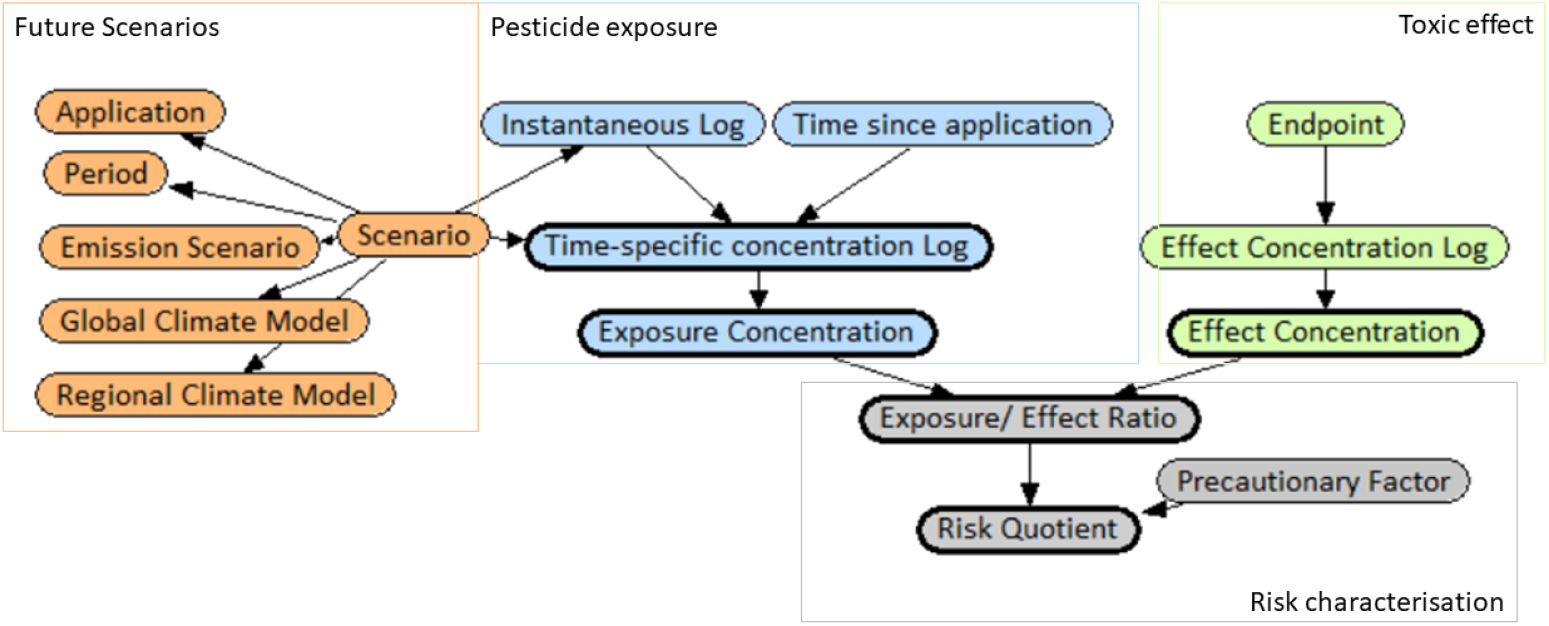
Conceptual model for the risk estimation of a pesticides. Pesticide exposure derives input from the WISPE platform and is determined by the associated Future scenarios. The Toxic effect nodes together with the exposure concentration derives the risk quotient under Risk characterization.

#### 2.1.2 Exposure sampling and modelling

Measured pesticide exposure concentrations, their distribution and associated uncertainties are highly influenced by sampling method, time and rate (Spycher et al., 2018). Data derived from monitoring has a wide range of uncertainties through sampling constraints and limited representativeness (FOCUS, 2007). Yet, a realistic environmental concentration is vital for good environmental risk assessment. This is especially significant whenever a single number is used without accounting for uncertainty, but is also influential when trying to derive a representative exposure distribution as uncertain estimations can hinder appropriate decision-making (Wolf and Tollefsen, 2021). Thence, the EU (Directive 2009/128/EC (EC, 2009a) and REGULATION (EC) No 1107/2009 (EC, 2009b)) offers the option to use models to predict environmental concentrations (PECs) in surface waters. Even if monitoring data are available, the use of modeling approaches for exposure assessment is encouraged by EFSA (2017). They have developed the FOCUS (FOrum for the Co-ordination of pesticide fate models and their Use) surface water scenarios using the model tool SWASH, a GUI for the models PRZM (Pesticide Root Zone Model), MACRO and TOXSWA (TOXic substances in Surface Waters). PRZM and MACRO are models frequently used to simulate pesticide transport in soil while TOXWA simulates the dilution at the edge of field or drain water concentration from the other two models in different surface water body types. SWASH takes agricultural management practices, climate, crops, topography, and soil types into account (Adriaanse et al., 2017). For this study we used the World Integrated System for Pesticide Exposure (WISPE) platform, that was developed to evaluate the potential for pesticide exposure to surface waters and groundwaters (Bolli et al., 2013). The WISPE platform was configured with scenarios containing crop, soil, and weather conditions for representative agricultural areas among others in the EU, USA and Norway. This modelling platform interlinks the pesticide root zone model (PRZM), an exposure analysis modeling system (EXAM) (Burns, 2004) and the aquifer dilution assessment model (ADAM) (Williams, 2010) similar to TOXWA. The PRZM model simulates the movement of chemicals within and below the root zone (in unsaturated soil systems). EXAM is a hydraulic model combined with a chemical fate and transport model simulating processes in aquatic environments. It simulates various processes in the aquatic environment. ADAM is an integrated model which predicts the chemical dilution, partitioning and persistence to a water body. EXAM and PRZM are standard models used by USEPA, and the latter model is also used in European pesticide registration and risk assessment (REGULATION (EC) No 1107/2009 (EC, 2009b)). In a previous study, the transport of particles and particle bound pesticides was calibrated for two field sites representative for Norwegian agricultural areas by Bolli et al. (2013). The study found that in this northern region the erosion and transport of particle bound pesticides are heavily dependent on the weather conditions such as precipitation shortly after application or melting-freezing episodes which take place in spring and winter.

### 2.2 Bayesian network modules

In the following, the information sources and assumptions for the four modules of the BN model and the model runs are described. The exposure model platform WISPE was run for several application scenarios and for two climate models. In the selected case study area, environmental factors such as soil and site parameters together with chemical properties and climate scenarios were linked to the exposure of a pesticide by using the WISPE platform. The probability distribution of pesticide exposure was obtained from predicted concentrations for multiple years, which enables accounting for variability over a longer time period (FOCUS, 2007). Correspondingly, for the probability distribution of effects, we used the range of species sensitivities from toxicity tests. The software Netica (Norsys Software Corp., www.norsys.com) was used to construct the BN. A more detailed node description and model assumptions are given in the following Table 1.

**Table 1.** Bayesian network node description detailing the type of node, the number of states and the equation used to train the network.

| Node name (Variable) | Type | States* | Netica equation and node input |
| --- | --- | --- | --- |
| Climate | Constant | 2 | Scenarios |
| Period | Constant | 3 | Scenarios |
| Application | Constant | 3 | Scenarios |
| Scenario | Constant | 18 | Combination of the scenarios |
| Intercept Log | DC ** | 5 | Probabilities for specific scenario |
| Time since application | Constant | 5 | WISPE model prediction for day 1, 2, 5, 21 OR 60 |
| Time specific concentration Log | DC | 10 | Scenario specific function:<br>Time specific concentration Log = Intercept Log + (slope *** x time)) |
| Endpoint | Constant | 2 | EC50 (day 1)<br>NOEC (day 1 – 61) |
| Effect concentration Log | DC | 10 | EC50: NormalDistribution (mean, sd) OR<br>NOEC: NormalDistribution (mean, sd) |
| Exposure concentration | DC | 10 | $\exp^{\text{Time specific concentration Log}}$ |
| Effect concentration | DC | 10 | $\exp^{\text{Effect concentration distribution Log}}$ |
| Exposure – effect – ratio | DC | 7 | Ratio - Exposure concentration distribution:<br>Effect concentration distribution |
| Precautionary factor | Constant | 5 | Either 1, 3, 10, 30, 100, 300, OR 1000 |
| Risk quotient | DC | 7 | Exposure – effect – ratio distribution x<br>Precautionary factor |

\* number of states
\*\*DC - discretized continuous; continuous variables were binned into the states
\*\*\*scenario depended average slope

#### 2.2.1 Future scenarios

The agricultural sector manages 3.5 % of Norway’s land area pr. 2021. Being part of the northern region, Norway has lower temperatures and a shorter growing season than central and southern Europe. These climate conditions restrict the area suitable for grain cultivation. Until the year 2060, the annual average temperature is expected to increase by approx. 2°C, with the largest increase in temperature in winter, and the lowest in summer in Norway. Consequently, the meteorological growing season will be longer than today’s, with a predicted increase in growing season of up to two months towards the end of the century (Fuglestvedt, 2016). This may lead to earlier sowing, ripening and harvest for spring cereals and growing of crop types that mature later but offer a higher yield potential. CC is also expected to lead to significant changes in precipitation with an increase of 8% for annual precipitation at the end of the century, but with large variation between the cropping regions in Norway. For the cultivation of grain, not only the amount and intensity of rainfall is of interest, also its frequency and distribution throughout the growing season. Other expected CC impacts are the introduction of new plant pathogens and pests from southern countries to northern areas while existing will be able to take advantage of a longer growing season and multiply faster than before. Also, changes in crop composition may lead to a change in the occurrence of the diseases and possibly new host-parasite interactions (Fuglestvedt, 2016). Furthermore, pesticides efficacy is affected by environmental factors such as temperature, precipitation and wind (Olesen and Bindi, 2002). In Norway, a longer growing season and more frequent pest infestations may require the use of more pesticides. A warmer climate is expected to result in increased production of winter wheat. The milder cold season may provide better overwintering conditions for plant pathogens, which might entail early and more severe infestation of the crop the following season. The most relevant measure apart from using resistant crop types is spraying of fungicides. In addition, early infestations require spraying both earlier and more frequently during the growing season (Fuglestvedt, 2016). Based on this winter wheat was chosen as the model crop for this study. A more detailed description of the expected CC is given by Hanssen-Bauer et al. (2015). In the following, the future scenarios used to run the WISPE platform are described.

##### 2.2.1.1 Climate scenarios

In this study, we used two sets of climate projections originally developed for the site Grue in the south east of Norway (ca 160 km North-east of Syverud/Ås) under the GENESIS project (2009-2014, https://cordis.europa.eu/project/id/226536). Both were derived from the greenhouse gas emission scenario “A1B”, which was developed to represent a future world of very rapid economic growth, low population growth and rapid introduction of new and more efficient technology, for a spatial resolution of 50 km. The two sets of climate projections were derived by two global climate models (GCM) which we will refer to as Climate Model 1 (C1) and Climate Model 2 (C2). The GCM of C1, “ECHAM5-r3”(Roeckner et al., 2004), was developed by the Max Planck Institute for Meteorology, and the GCM for C2, “HADCM3-Q0”(Gordon et al., 2000), was developed at the Hadley Centre. Regional climate models (RCMs) are commonly applied to downscale from the global to more local levels (Jones et al., 2011, Samuelsson et al., 2011). Here, the same RCM called RCA3 was used, developed by the Rossby Center at SMHI (the Swedish Meteorological and Hydrological Institute). Thereby, C1 represents the regional climate model “ECHAM5-r3 A1B-SMHI-RCA3” and C2 represents “HADCM3-Q0 A1B-SMHI-RCA3”.

The climate projections used in this study has several limitations: the emission scenario and the two climate models are rather old, and they have not been bias-corrected for the study area. Moreover, climate projections should ideally be obtained from a larger ensemble of climate models rather than one or a few models. However, generating a new and more appropriate set of climate projections was beyond the scope of this study. Therefore, the climate projections that were already derived for the WISPE platform were considered sufficient for the purpose of demonstrating this BN approach.

Projections from the two climate models (C1 and C2) differed in precipitation, temperature, evapotranspiration, solar radiation and wind. For example, they had different projected changes in number of days with snow cover and changes of annual rainfall (Kjellstöm et al., 2011). The differences between the two climate models are especially of interest for the chosen days and months of pesticides application. Based on Mann Kendall (MK) trend analysis, C1 showed a positive trend in temperature, evapotranspiration and precipitation for a 3-day average before the day of pesticide application. When comparing climate conditions for 10-day average before day 21 after application, a positive trend could be detected for temperature and evapotranspiration (i.e. the process of water evaporation from soil and other surfaces through transpiration from plants). In general, C2 showed no trend, and even a negative trend for October for a 3-day average before the day of application (see Supplement Information I -Table S.2). The projections from C1 were more consistent with more recent climate projections for Norway, which show that an increase in temperature and precipitation can be expected (Hanssen-Bauer et al., 2015). Consequently, in this paper we decided to focus mainly on predicted exposure concentration based on C1.

##### 2.2.1.2 Pesticide application scenarios

The first pesticide application scenario is based on the current “common practice” dosage and referred to as the baseline (see Table 2). The second scenario is related to the European Green Deal, which aim for a 50% reduction of the pesticide use by 2030 (EC, 2020) (referred to as baseline-50%). The third scenario represent a potentially increased use of pesticides in the future, for example due to changing climate conditions and increased pest pressures (Fuglestvedt, 2016) (referred to as baseline+50%).

**Table 2.** Description of application scenarios used in this case study for the five selected pesticides

| Active substance | Unit | Dose active substance (g/daa) | Baseline - 50% | Baseline | Baseline +50% |
| --- | --- | --- | --- | --- | --- |
|  |  |  | Dose active substance (kg/ha) | Dose active substance (kg/ha) | Dose active substance (kg/ha) |
| **Clopyralid** | mL/daa | 5 | 0.025 | 0.05 | 0.075 |
| **Fluroxypyr-meptyl** | mL/daa | 10 | 0.05 | 0.1 | 0.15 |
| **MCPA** | mL/daa | 50 | 0.25 | 0.5 | 0.75 |
| **Trifloxystrobin** | mL/daa | 15 | 0.075 | 0.15 | 0.225 |
| **Prothioconazole** | mL/daa | 17.5 | 0.0875 | 0.175 | 0.2625 |

We selected active pesticide ingredients that are all approved in Norway for crop protection in winter wheat. Two plant protection products, a herbicide containing MCPA (CAS nr. 94-74-6), fluroxypyr-meptyl (CAS nr. 81406-37-3) and clopyralid (CAS nr. 1702-17-6), and a fungicide composed of trifloxystrobin (CAS nr. 141517-21-7) and prothioconazole (CAS nr. 178928-70-6), were chosen for the purpose of demonstrating the approach. Inherent properties such as molecular weight, water solubility, sorption properties (Koc), degradation half-life (DT50 soil), and vapor pressure, and Freundlich exponent (1/n), and systemic property e.g. plant uptake factor, were collected and included in the data asset (see Supplement Information I - Table S.1).

The associated recommended application rate and time of spraying were used to define the application scenario for the WISPE platform runs. It was assumed that the herbicide is applied once in the first half of May (crop growth stage BBCH 13-14; cf. label for Ariane ™ S, Corteva Agriscience), and that the fungicide is applied once in the first half of October (after sowing and germination of the winter wheat; cf. label for Delaro SC 325, Bayer Crop Science). For the calibration of the WISPE platform no tillage was assumed. Some of the combinations chosen for pesticide application, e.g. the choice of no soil tilling in combination with winter cereals, may not be the most common/optimal agronomic practice and can hence add to some of the uncertainty in the modelling.

#### 2.2.2 Pesticide exposure

The scenarios described above were used as input information for the WISPE platform. Additional settings used to run the platform are described in the succeeding section. The exposure distributions used as input for the pesticide exposure module were based on the predicted exposure concentration from the WISPE platform.

##### 2.2.2.1 WISPE platform settings

When the WISPE platform was first developed as a tool to estimate pesticide exposure in ground- and surface water for Norwegian conditions, two study areas were chosen as representative field sites to generate data for calibration and validation of the model (Bolli et al., 2013). In this study, we used the Syverud site scenario, which was developed to represent larger agricultural areas in South East Norway. The study site is located on the grounds of the Norwegian University of Life Sciences (NMBU) in Ås (Supplement Information I – Figure S. 1). The soil in this study area is classified as loam/ silt loam, with 26% clay, 49% silt and 25% sand content. The area was formerly used as a meadow which resulted in a soil structure with high infiltration capacity, aggregate stability and saturated hydraulic conductivity (Bolli et al., 2013). For the model simulations the site was assumed to be ploughed in autumn, with a ploughing depth of 20 cm. The platform predicts output concentrations for a stream, pond and ditch with parameters adapted originally from TOXSWA into the EXAM model. We have only considered the predicted output for the stream environment, with the following water body parameters: 1 m width, 100 m total length, 0.3 m average water depth, 15 mg/L concentration of suspended solids, 5 % organic carbon content, and 800 kg/m^3^ dry bulk density (FOCUS, 2015). WISPE was calibrated for the model crop winter wheat.

##### 2.2.2.2 Exposure prediction platform implementation

The WISPE platform is run according to the previously mentioned future scenarios and platform settings such as the selected representative field site, crop type and for the various time-periods of C1 and C2. The WISPE platform predicts exposure concentration for 26 years, corresponding to the 26 years over which the model runs. The concentrations are predicted for instantaneous, 24 h, 96 h, 21, 60 and 90 days. In the further process, the time-periods were changed into periods of 30 years (2000-30, 2035-65, 2070-00) to derive the distributions (BN input). The platform simulated pesticides to specified unique application conditions for the two climate models. In total, 18 scenarios were used in the developed BN per pesticide (Table 3).

**Table 3.**
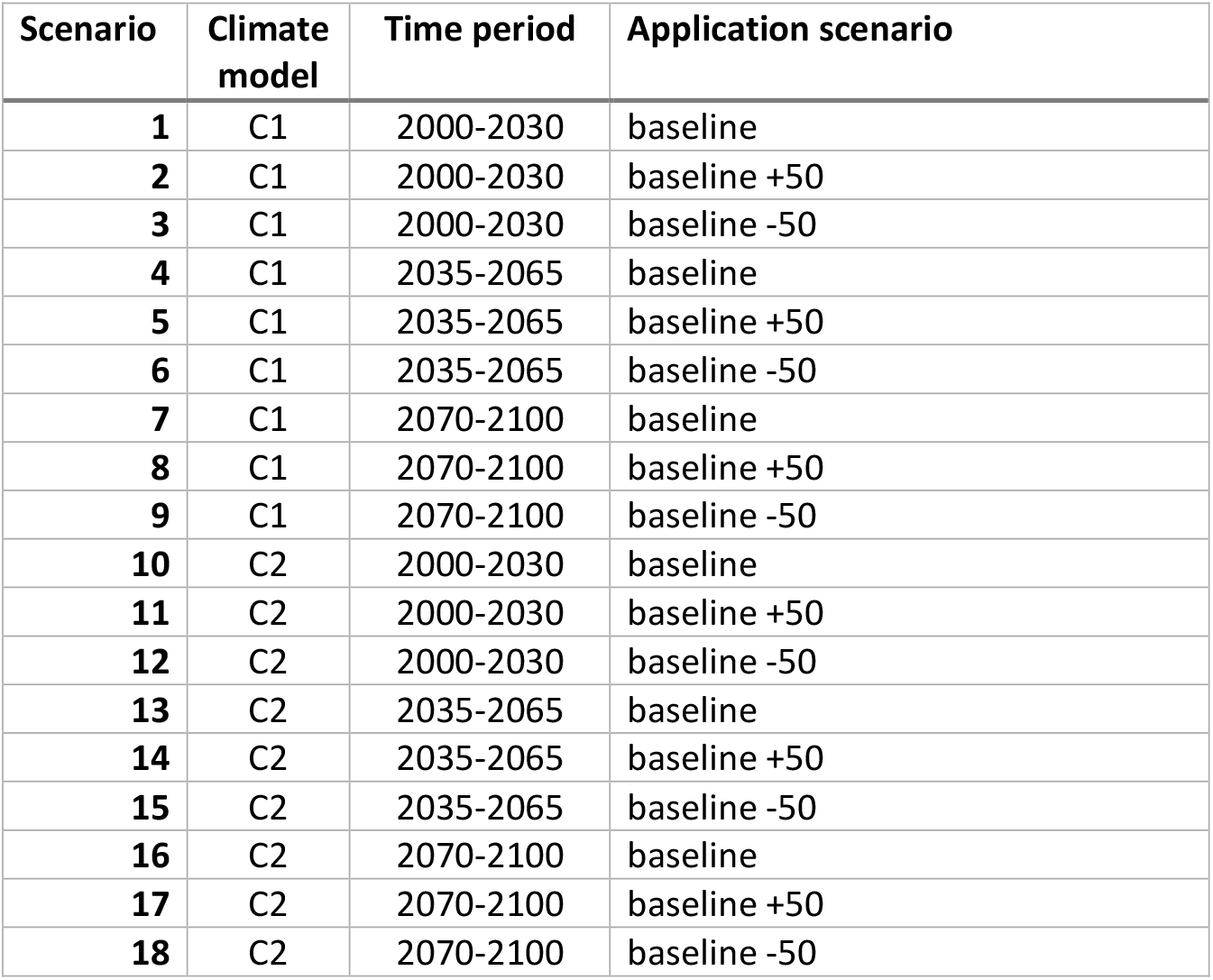
Scenario overview used in the developed Bayesian network model to derive climate, time period and application specific risk quotient distribution

| Scenario | Climate model | Time period | Application scenario |
| --- | --- | --- | --- |
| 1 | C1 | 2000-2030 | baseline |
| 2 | C1 | 2000-2030 | baseline +50 |
| 3 | C1 | 2000-2030 | baseline -50 |
| 4 | C1 | 2035-2065 | baseline |
| 5 | C1 | 2035-2065 | baseline +50 |
| 6 | C1 | 2035-2065 | baseline -50 |
| 7 | C1 | 2070-2100 | baseline |
| 8 | C1 | 2070-2100 | baseline +50 |
| 9 | C1 | 2070-2100 | baseline -50 |
| 10 | C2 | 2000-2030 | baseline |
| 11 | C2 | 2000-2030 | baseline +50 |
| 12 | C2 | 2000-2030 | baseline -50 |
| 13 | C2 | 2035-2065 | baseline |
| 14 | C2 | 2035-2065 | baseline +50 |
| 15 | C2 | 2035-2065 | baseline -50 |
| 16 | C2 | 2070-2100 | baseline |
| 17 | C2 | 2070-2100 | baseline +50 |
| 18 | C2 | 2070-2100 | baseline -50 |

The following example shows the log transformed WISPE output for scenario 11 (Figure 2). A log-linear equation was fitted to the time series concentrations generated by the WISPE platform using R (R version 4.1.0), the tidyverse package (Wickham et al., 2019) and some base R functions (Team, 2020). The average slope and the probabilities for each scenario were generated (see Supplement Information II) and used as input in the conditional probability table (CPT) of the instantaneous concentration and time specific concentration node (see Supplement Information II).

**Figure 2.**
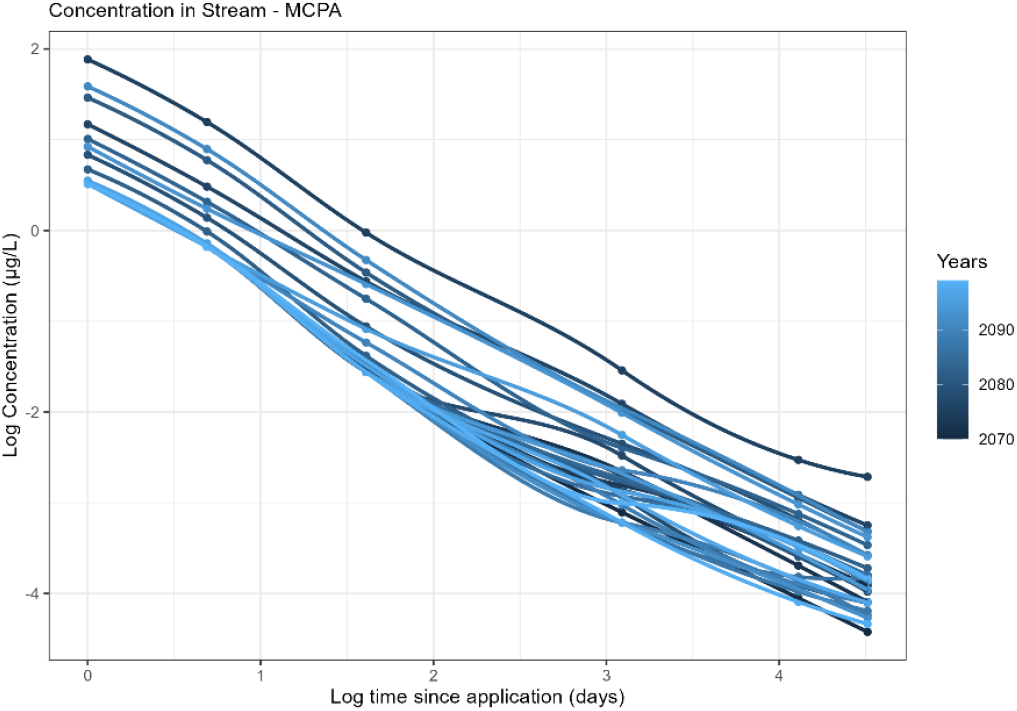
Example log-linear regression of the concentration in the stream predicted by the WISPE platform for MCPA with an application baseline scenario for the Climate model 1 and the time interval 2070-2100.

In general, the instantaneous node interval range differs for each selected pesticide: clopyralid 0.0025 to 0.6065 μg/L, fluroxypyr-meptyl 0.0111 to 4.4817 μg/L, MCPA 0.0821 to 12.1825 μg/L, prothioconazole 0.0302 to 0.2231 μg/L, trifloxystrobin 0.0235 to 0.1653 μg/L. Taking a closer look at precipitation, one of the determining climate conditions for the transport and fate of pesticides, it can be observed that with higher amount of precipitation the exposure concentration increase (Figure 3). In addition, it can be noted that there is an interaction between application and precipitation as the effect of amount of precipitation is higher (steeper slope) when the pesticide application is higher to start with (baseline+50%).

**Figure 3.**
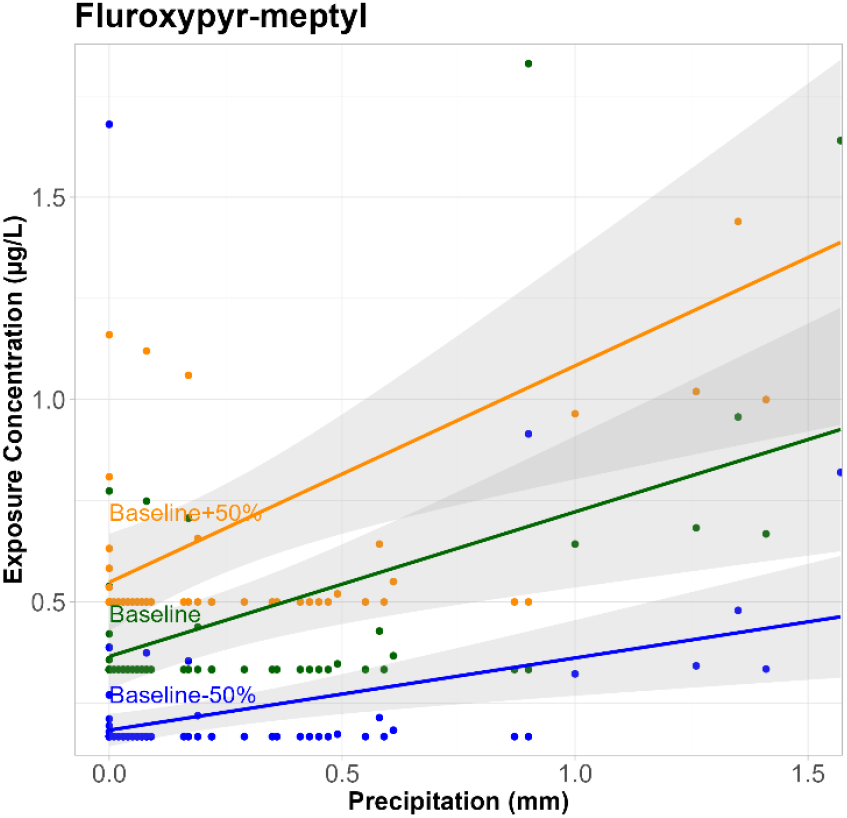
Example fluroxypyr-meptyl exposure concentration vs precipitation for the three tested application scenarios for a 3-day sum of precipitation before the day of application (here: 5-7 October for the period 2000-2100).

#### 2.2.3 Pesticide effects

Uncertainties related to current effect assessment are often associated with extrapolation from laboratory to field and inter-intraspecies variation (Rai et al., 2002) and can also be linked to the data set size. In traditional regulatory effect assessment these uncertainties are usually accounted for by assessment factors. In this study, NOEC values (no-observed effect concentrations) and EC50 values (effect concentration for 50% of the test population) were used as input in the pesticide effect module. The effect distributions were fitted to toxicity data for NOEC (and NOEL) or EC50 (and LC50) endpoints. The applied probabilistic model is similar to a species sensitivity distribution (SSD), for variation in sensitivity of species to toxicants which is used extensively in ecotoxicology (Belanger and Carr, 2020). SSDs are now commonly used as an alternative to the conservative approach on the basis of the most sensitive species (lowest NOEC value). They are based on multiple toxicity tests of different species and thereby reflect interspecies differences in sensitivity to a chemical. Subsequently, SSDs can be used to develop a community level threshold (Belanger et al., 2017). Modelling and data requirements guidance was found in the “Technical Guidance for Deriving Environmental Quality Standards” (TGD) (SCHEER, 2017). We mainly used toxicity data from the US EPA ECOTOX Knowledgebase (https://cfpub.epa.gov/ecotox/search.cfm) and supplemented this with data from Middle Tennessee State University EnviroTox Database (https://envirotoxdatabase.org). The EC50 effect distribution was derived from EC50 and LC50 toxicity data (Table 4). The NOEC distribution is based on NOEC and NOEL values, apart from Clopyralid for which only NOEC toxicity data was available. If multiple values for the same species occurred in the data set, the mean was used as a data point to derive the distribution (Mentzel et al., 2021). The number of observations for this study varied depending on the chemical, and whether it was an EC50 or NOEC toxicity test. In our research, we only considered adverse effects such as mortality, reproduction and growth. The distribution was derived using the R package *MASS* (Venables and Ripley, 2002). The data preparation was carried out with the R package *tidyverse* (Wickham et al., 2019) (see Supplement Information III)

**Table 4.** Effect (toxicity) data collected for this study, detailing the effect groups per pesticides and the derived EC50 and NOEC natural log (ln) mean and standard deviation for the natural log distributions.

|  | Number of values |  | Effect | EC50 |  | NOEC |  |
| --- | --- | --- | --- | --- | --- | --- | --- |
|  | EC50 | NOEC |  | Ln Mean (µg/L) | Ln sd | Ln Mean (µg/L) | Ln sd |
| **Clopyralid** | 7 | 8 | Growth, Population, Reproduction, Development, Mortality | 11.45 | 1.99 | 7.73 | 3.67 |
| **Fluroxypyr-meptyl** | 16 | 11 | Population, Mortality | 7.08 | 2.06 | 6.03 | 2.01 |
| **MCPA** | 45 | 20 | Population, Mortality, Growth, Morphology, Development, Reproduction, | 9.56 | 3.11 | 7.07 | 2.10 |
| **Trifloxystrobin** | 19 | 17 | Growth, Development, Mortality, Population, Morphology | 4.48 | 1.72 | 3.18 | 1.77 |
| **Prothioconazole** | 11 | 10 | Population, Mortality, Growth | 7.41 | 1.78 | 6.17 | 2.00 |

#### 2.2.4 Risk characterization

This module consists of three nodes: exposure/effect ratio, a precautionary factor and the risk quotient (RQ). The assumptions for the node input are described in Table 1. The BN was run for the different scenarios and with either an EC50 (and day 1 since application) or a NOEC (and day 1-61 since application) distribution. As explained in Mentzel et al. (2021), the precautionary factors have a similar role as the assessment factors that are frequently used in risk estimation (see TGD (SCHEER, 2017)). A risk is assumed when the RQ is higher than 1 (Bruijn et al., 2002). Following the findings in Mentzel et al. (2021), all RQ distributions displayed in the results were derived with a precautionary factor of 10 (see Figure 4). A higher precautionary factor will increase the probability of the RQ exceeding 1. The output for each of the settings (evidence) was collected and can be found in the Supplement Information IV. It contains a detailed collection of the probabilities for each of the RQ node intervals depending on the selected evidence. We carried out a MK trend analysis for the predicted exposure concentration (WISPE platform output) for baseline application, C1 and C2, Day 1 and 21 since application and for all of the selected pesticides (see Table 5). A positive trend indicates an increase of the predicted exposure concentration. The trend was assumed to be negative whenever τ < 0 and p < 0.1. The trend was assumed positive when τ > 0 and p < 0.1.

**Table 5.** MannKendall trend analysis for the predicted pesticide exposure of selected pesticides for baseline application in May and October for acute and chronic conditions for climate model 1 (C1) and climate model 2 for (C2) for the whole period of time 2000-2100

| Scenario | Application | Climate model | Days since application | Pesticide | Type | Time of application | $\tau$ | p |
| --- | --- | --- | --- | --- | --- | --- | --- | --- |
| 1 | baseline | C1 | 1 | Clopyralid | Herbicide | May | -0.005 | 0.951 |
| 1 | baseline | C1 | 1 | Fluroxopyr- meptyl | Herbicide | May | 0.038 | 0.634 |
| 1 | baseline | C1 | 1 | MCPA | Herbicide | May | 0.034 | 0.655 |
| 1 | baseline | C1 | 1 | Prothiocanazole | Fungicide | October | **0.138** | **0.084** |
| 1 | baseline | C1 | 1 | Trifloxystrobin | Fungicide | October | **0.182** | **0.024** |
| 1 | baseline | C1 | 21 | Clopyralid | Herbicide | May | 0.006 | 0.935 |
| 1 | baseline | C1 | 21 | Fluroxopyr- meptyl | Herbicide | May | -0.020 | 0.773 |
| 1 | baseline | C1 | 21 | MCPA | Herbicide | May | 0.048 | 0.490 |
| 1 | baseline | C1 | 21 | Prothiocanazole | Fungicide | October | 0.039 | 0.618 |
| 1 | baseline | C1 | 21 | Trifloxystrobin | Fungicide | October | 0.082 | 0.312 |
| 10 | baseline | C2 | 1 | Clopyralid | Herbicide | May | 0.111 | 0.142 |
| 10 | baseline | C2 | 1 | Fluroxopyr- meptyl | Herbicide | May | **0.159** | **0.044** |
| 10 | baseline | C2 | 1 | MCPA | Herbicide | May | **0.133** | **0.078** |
| 10 | baseline | C2 | 1 | Prothiocanazole | Fungicide | October | 0.002 | 0.983 |
| 10 | baseline | C2 | 1 | Trifloxystrobin | Fungicide | October | -0.054 | 0.506 |
| 10 | baseline | C2 | 21 | Clopyralid | Herbicide | May | 0.034 | 0.620 |
| 10 | baseline | C2 | 21 | Fluroxopyr- meptyl | Herbicide | May | 0.078 | 0.269 |
| 10 | baseline | C2 | 21 | MCPA | Herbicide | May | 0.054 | 0.432 |
| 10 | baseline | C2 | 21 | Prothiocanazole | Fungicide | October | -0.125 | 0.111 |
| 10 | baseline | C2 | 21 | Trifloxystrobin | Fungicide | October | **-0.14** | **0.080** |

**Figure 4.**
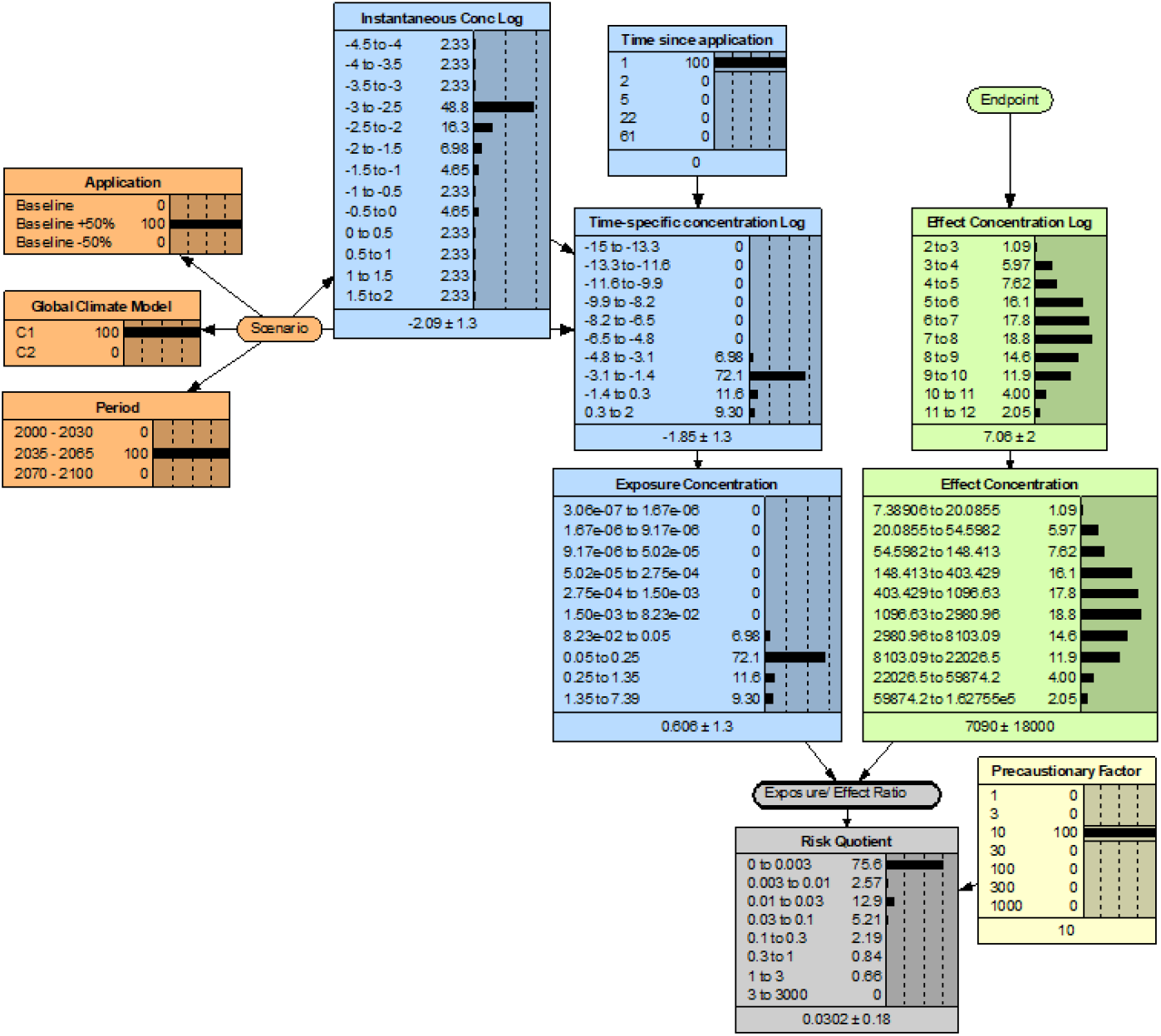
Example of the Bayesian network parameterized for fluroxypyr-meptyl, with a baseline+50% application, global climate model C1, time period of 2035-2056, for a time since application of 1 day and a EC50 based effect distribution.

## 3 Results

### 3.1 Predicted pesticide exposure

The pesticide concentrations predicted by the WISPE platform predictions showed similar temporal trends as the those described for the climate variables (see Supplement Information I -Table S. 2). The Mann-Kendall trend analysis showed mostly no trends over the whole range of years (2000-2100), for the different pesticides and seasons. However, some of the trends we observed for the climate variables were reflected in the exposure prediction, such as the positive trend in fungicide exposure in October for C1. Also, there was a negative trend derived for trifloxystrobin and a positive for fluroxypyr-meptyl and MCPA for C2 (Table 5).

### 3.2 Risk distribution for the time after application

In the following, the RQ node outputs are described in a comprised form visualizing the results for the different scenarios (e.g. Table 3 for reference) and are displayed in a stacked bar for easy comparison. The RQ interval range was from 0 to 3000 or 20000 (MCPA), with 0 being no risk (blue) and 3-3000 or 20000 being high risk (red). The RQ was either derived with from NOEC values (RQ_NOEC_) or from EC50 values (RQ_EC50_) for a baseline application. This analysis enabled the identification of periods with higher risk of environmental effects of individual pesticides or groups of pesticides.

The RQ_NOEC_ distribution interval range and their probabilities varied for each of the selected pesticides. As expected, the RQ_EC50_ distribution generally showed higher probabilities of being in the lowest RQ interval “0-0.003” compared to the RQ_NOEC_ (at day 1 after application). For clopyralid applied in May, the RQ_NOEC_ was most likely (80% probability) in the lowest interval (see Supplement Information I - Figure S. 3). For the fungicide prothioconazole, applied in October, the probability of RQ_NOEC_ being below 0.01 was 90%. The RQ of MCPA had the widest (range) of distribution, with about 10% probability of the RQ_NOEC_ exceeding 0.1. For one day since application, this herbicide also had a probability of about 1 % of being the highest RQ interval ‘3-20000’ (dark red) (Figure 5a). The fungicide trifloxystrobin had the lowest probability of RQ_NOEC_ being in the lowest interval of ‘0-0.003’, compared to the other pesticides for the same day since application (Figure 5b). Clopyralid and trifloxystrobin both had an interval range from 0 to 1, fluroxypyr-meptyl ranged to 3, and prothioconazole ranged to 0.3 (see Supplement Information II). MCPA had higher probability of the RQ_NOEC_ being below 0.01 whereas trifloxystrobin had higher probability of being below 0.1.

**Figure 5.**
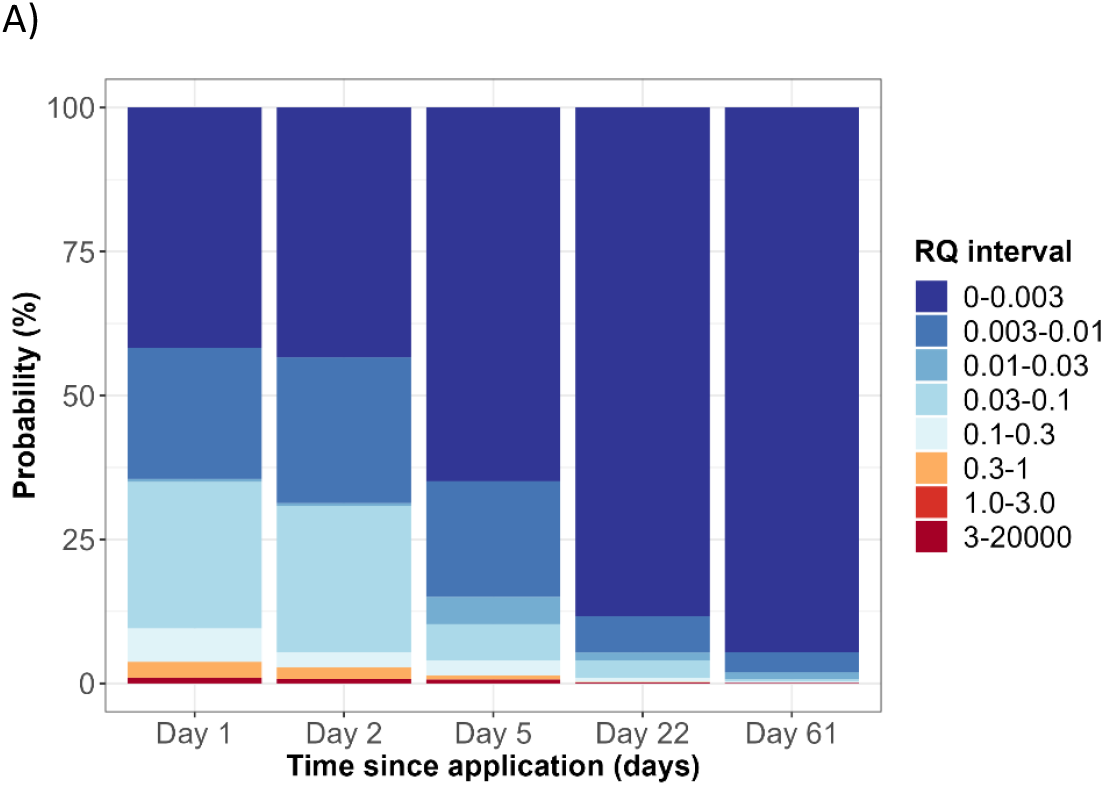
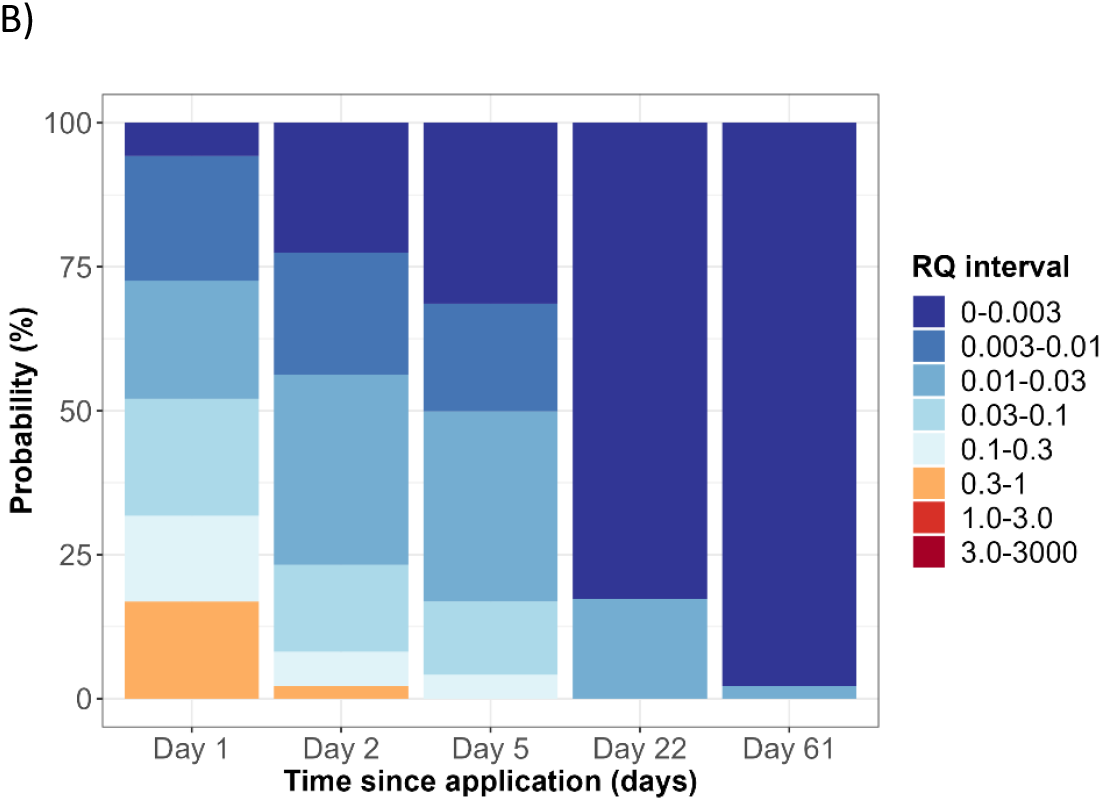
Example of MCPA (A) and trifloxystrobin (B) risk estimation over time for 1, 2, 5, 22, and 61 days after application, for the baseline application scenario, climate model C1 and the time interval of 2070-2100 for NOEC based effect distribution.

### 3.3 Plausible scenarios: combination of climate change and pesticide application

A change in pesticide application patterns such as an increase in the rates or number of applications per season, can be considered as adaptation to consequences climate change (e.g. increased pest pressure). We therefore consider scenario combining future climate projections (period 2035-2065) with increased pesticide application as a plausible scenario. On the other hand, the combination of future climate projections with reduced pesticide application represent a scenario more in line with EU’s pesticide policy. Therefore, we compare the RQ_EC50_ of the current time period (2000-2030) and baseline application with the predicted RQ_EC50_ for a future time period (2035-2065) and both baseline-50% and baseline+50% application scenarios. The figures show examples for the fungicide trifloxystrobin (Figure 6a) and the herbicide fluroxypyr-meptyl (Figure 6b). Trifloxystrobin had more than 10% probability of RQ_EC50_ exceeding 0.03 for the current practice. In future, applying less fungicide will result in a shift towards lower RQ intervals and an overall decrease in risk. From the BN predictions we also observed the general tendency that applying 50% less resulted in a shift towards lower RQ intervals, whereas applying 50% more only resulted in a minimal shift towards higher RQ intervals. This was also the case for some of the other pesticides, which appeared to have more similar RQ_EC50_ distributions for baseline and baseline+50% (see Supplement Information I - Figure S. 4). Fluroxypyr-meptyl, on the other hand, showed a clear trend towards higher RQ_EC50_ states for the baseline+50% application. In general, the probability of RQ_EC50_ exceeding 1, which commonly used as a threshold for concern, was low and not much influenced by the different time periods or application scenarios.

**Figure 6.**
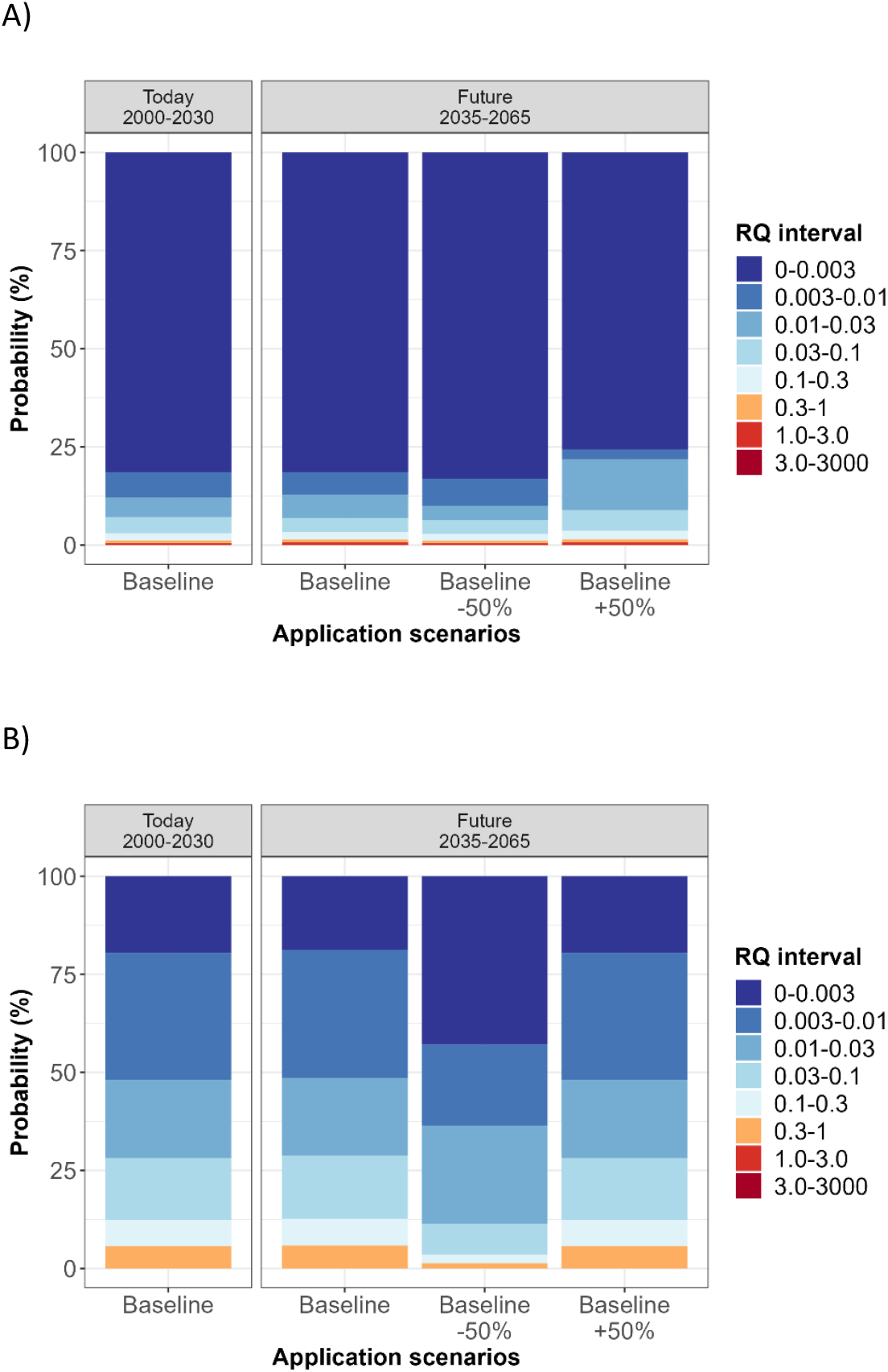
Risk estimation of a selected herbicide and fungicide, for a time since application of 1 day, for the climate model C1 and for EC50 based effects distribution. The herbicide, fluroxypyr-meptyl (A), displays scenarios 1,4,6 and the fungicide, trifloxystrobin (B), displays the scenario 1,7,8 Table 3.

## 4 Discussion

As monitoring of environmental pesticide concentration is costly and time-consuming, future climate conditions need to be incorporated better in risk assessment. The complexity and lack of well characterized processes in pesticide risk assessment can be overcome by taking advantage of the BNs’ ability to use data from various different sources, which is one of their benefits (Hamilton and Pollino, 2012, Troldborg et al., 2021, Mentzel et al., 2021, Gibert et al., 2018). Moreover, they can be constructed as causal models that help comprehend hazard pathways and vulnerability relations better and with that assist in risk prioritization (Sperotto et al., 2017). A study by Gaasland-Tatro (2016) showed how CC factors can be included in BNs by using a relative risk model that evaluates ecological parameters over landscape scale regions and other stressors. Along these lines, Landis et al. (2013) pointed out that today’s environmental risk assessment should also consider interactions among contaminant and noncontaminant stressors, together with new regimes of precipitation and temperature at specific geographical sites (Landis et al., 2013). An example for the use of multiple information sources as inputs for a BN has also been presented by Troldborg et al. (2021). They used a causal network developed for pesticide risk analysis for a drinking water catchment that was informed by expert knowledge and used GIS for spatial risk assessment. The use of BN models in ecotoxicology is still rare compared to other types of environmental assessment (Kaikkonen et al., 2021) even though their use has increased in chemical risk assessment in recent years (Moe et al., 2021a). One of the common shortcomings of BNs is the loss of precision due to discretization of continuous variables; this phenomenon was also observed for some of the selected pesticides in our study, e.g. MCPA. Although the instantaneous pesticide concentration distribution differed between the baseline and baseline+50% scenarios, these differences were not reflected in the exposure concentration node, where the probability distribution appeared very similar. This resulted in similar RQ distribution for the two application scenarios, and can lead to loss of information due to the discretization of continuous variables (Marcot, 2017, Nojavan et al., 2017). This technical problem can be amended by through dynamic discretization which can enable higher resolution and lower uncertainty of the BN predictions (Carriger et al., 2016, Fenton and Neil, 2018).

The applicability domain of the BN model presented here is constrained by the current applications and calibration of the WISPE model platform. Until now, the WISPE platform was validated by Bolli et al. (2013) and offers the possibility to predict environmental concentrations for specific and representative study fields in Norway. The platform takes into account chemical properties and environmental factors when predicting the exposure of pesticides in the selected water body (Bolli et al., 2013). A predicted exposure time series with multiple peak concentrations could not easily have been incorporated in the exposure module of the BN, which currently assumes a log-linear decrease in pesticide concentration over time. Further development of this module would be needed to account for a more complex temporal exposure pattern.

The BN model we have developed based on the used input data predicts a slight increase in the probability of RQ exceeding 1 for future time periods. In other words, the model predicts higher risk for aquatic organisms with the climate projection data for the intermediate and last time periods investigated. This is expected and consistent with what has previously suggested regarding pesticide fate and transport being influenced by precipitation in northern Europe: An increased precipitation in future can imply increase risk of pesticides to freshwater ecosystems in agricultural areas.

As mentioned, the climate models used in this study were not properly bias-corrected for the study area. Thus, improved model precision and realism could be achieved by using more updated climate projections based on a larger number of climate models. In our opinion, further model development with a newer and refined version of the WISPE, could reduce some of the uncertainty related to predictions. Some beneficial improvements could be made in the structure of input and output files, as well as an interface easily connectable to other “programming tools” enabling fast and easy run for various differing input scenarios.

In addition, extending the constructed BN with more applications, updated and site-specific climate scenarios, crop and pesticide types, and the use of other representative study areas would be beneficial for the integration of variability in model predictions. This BN model could also be further developed to predicting the cumulative risk of intentional pesticide mixtures.

## 5 Conclusion

With this study, we demonstrate how exposure prediction model inputs and outputs can be incorporated into a BN deriving a RQ distribution for various events (scenarios). The constructed network propagates and incorporates uncertainty of all components in a transparent way when carrying out the probabilistic risk estimation. In principle, the BN model predicted a slight increase in the probability of RQ exceeding 1 for the intermediate (2035-2065) and latest time period (2070-2100) compared to the current (2000-2030). Further plans for this BN are to integrate more updated and bias-corrected climate projections from a larger ensemble of climate models. Our future research efforts will also explore more advanced options for risk characterization as alternatives to the currently used RQ approach, for example to make better use of causal dose-response relationships in cases where such information can be obtained. Future research should therefore consider approaches that incorporate not only an exposure prediction model under alternative future conditions but also an effect prediction model. Nevertheless, the presented approach shows promise in its ability to characterize the environmental risk of pesticides under future scenarios by integrating different types of information from agricultural practice, climate models, pesticide exposure models and toxicity testing.

## Acknowledgement

We thank Randi Bolli, (NIBIO), discussion and advice on the pesticide exposure modelling, as well as Wayne Landis (Western Washington University) and John Carriger (USEPA) for advice on Bayesian network modelling.

## Software availability

A free version of NeticaTM is available online at: http://www.norsys.com/downloads.html, along with a glossary of BN terms and tutorials (though the free version has a limited number of nodes and other limitations in functions). The WISPE model is available upon request for more information: https://www.nibio.no/prosjekter/oppdatering-av-modellen-wispe-og-de-norske-overflatevannscenariene

