## Supplement Information I for "Probabilistic risk assessment of pesticides under future agricultural and climate scenarios using a Bayesian network"

1. **Study area**

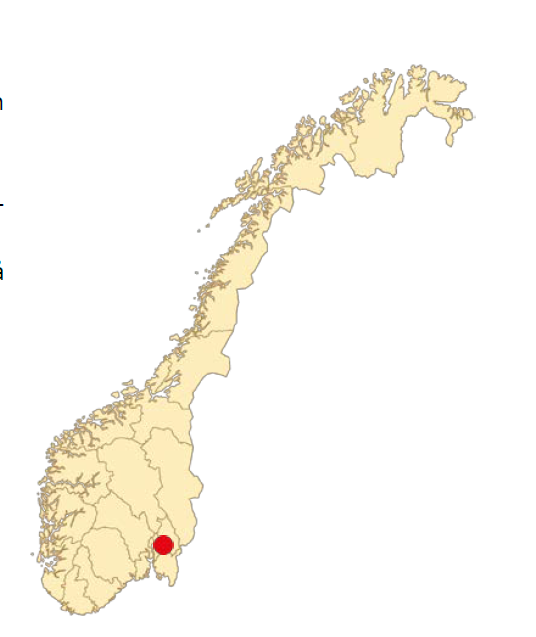

Figure S. 1 Map of Norway detailing the Syverud location (red dot).

2. **Pesticide properties**

Table S. 1 Summary of pesticide properties considered in the WISPE platform (Lewis et al., 2016)

|  | Mol. Weight | Solubility, mg/L | Plant Uptake Factor | Koc | DT50soil, lab, days | Vapour pressure, mPa | Freundlich exp. 1/n | EXAMS Aerobic metabolism, days (DT50water) | Q10, 20 °C | EXAMS Anaerobic met. days (DT50 sediment) | Direct Photolysis, days |
| --- | --- | --- | --- | --- | --- | --- | --- | --- | --- | --- | --- |
| Clopyralid | 192 | 7850 | 0.5 | 5 | 23.2 | 1.36E-09 | na * | 148 | 2.58 | 1000 | 271 |
| Fluroxypyr-meptyl | 367.24 | 0.136 | 0 | 19550 | 1 | 0.01 | na * | 34.7 | 2.58 | 1000 | 63 |
| Mcpa | 200.62 | 29390 | 0.5 | 74 | 24 | 0.4 | 0.68 | 13.5 | 2.58 | 1000 | 0.05 |
| Prothioconazole | 344.26 | 22.5 | 0.5 | 2556 | 0.44 | 7.40E-06 | 0.88 | 0.0 | 2.58 | 1000 | 2.1 |
| Trifloxystrobin | 408.37 | 0.61 | 0 | 2287 | 0.34 | 3.40E-03 | 0.96 | 1.1 | 2.58 | 1000 | 2.7 |

3. **Comparison Climate variables Climate model 1 & 2: Mann-Kendall -trend analysis**

Table S. 2 Mann-Kendall trend analysis for Climate variables of Climate model 1 (C1) and Climate model 2 (C2) for mean day since application (mean over 3 previous days) and 21 days (mean of 10 previous days) climate conditions for application in May and October.

|  |  | May |  |  | October |  |  |
| --- | --- | --- | --- | --- | --- | --- | --- |
| Days since application | output | MK.tau | MK.p | conclusion | MK.tau | MK.p | conclusion |
| 1 | MeanTemp1 | 0.167 | 0.252 | zero | **0.273** | **0.059** | pos |
| 1 | MeanPrecip1 | -0.011 | 0.962 | zero | **0.361** | **0.012** | pos |
| 1 | MeanEpot1 | **0.244** | **0.093** | pos | **0.25** | **0.084** | pos |
| 1 | MeanWind1 | 0.196 | 0.191 | zero | 0.131 | 0.374 | zero |
| 1 | MeanRadiat1 | -0.067 | 0.657 | zero | 0 | 1 | zero |
| 1 | MeanTemp2 | 0.16 | 0.272 | zero | 0.053 | 0.726 | zero |
| 1 | MeanPrecip2 | 0.135 | 0.369 | zero | **-0.32** | **0.029** | neg |
| 1 | MeanEpot2 | 0.05 | 0.744 | zero | -0.127 | 0.387 | zero |
| 1 | MeanWind2 | -0.168 | 0.264 | zero | 0.172 | 0.242 | zero |
| 1 | MeanRadiat2 | **-0.36** | **0.012** | neg | -0.067 | 0.657 | zero |
| 21 | MeanTemp1 | **0.249** | **0** | pos | **0.301** | **0** | pos |
| 21 | MeanPrecip1 | -0.029 | 0.676 | zero | 0.015 | 0.83 | zero |
| 21 | MeanEpot1 | **0.279** | **0** | pos | **0.318** | **0** | pos |
| 21 | MeanWind1 | 0.018 | 0.79 | zero | -0.037 | 0.588 | zero |
| 21 | MeanRadiat1 | -0.063 | 0.358 | zero | **-0.123** | **0.072** | neg |
| 21 | MeanTemp2 | **0.267** | **0** | pos | **0.451** | **0** | pos |
| 21 | MeanPrecip2 | **-0.124** | **0.069** | neg | 0.088 | 0.201 | zero |
| 21 | MeanEpot2 | **0.196** | **0.004** | pos | **0.412** | **0** | pos |
| 21 | MeanWind2 | 0.072 | 0.29 | zero | 0.07 | 0.308 | zero |
| 21 | MeanRadiat2 | -0.005 | 0.942 | zero | **-0.195** | **0.004** | neg |

**4 Visualized Result output for all selected pesticides for direct and indirect climate effect**

4.1 **Time since application**

a)

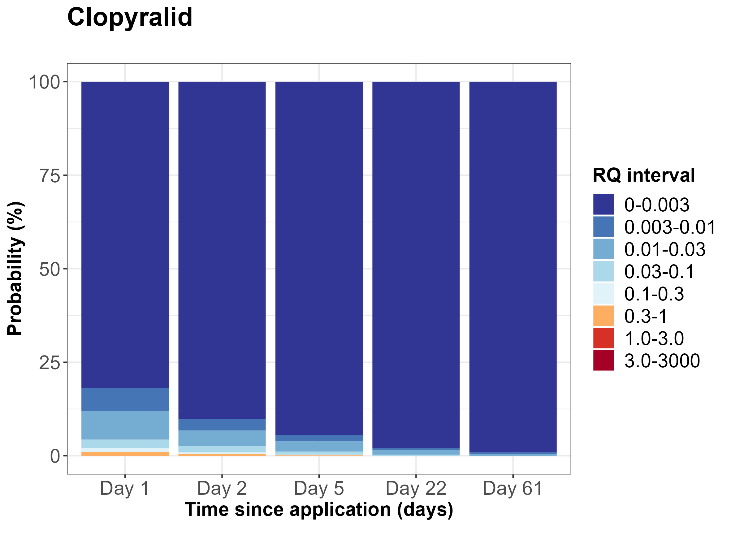

b)

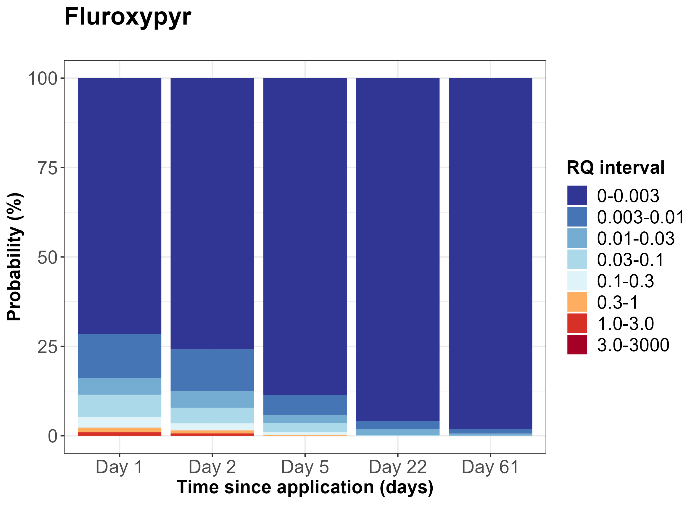

c)

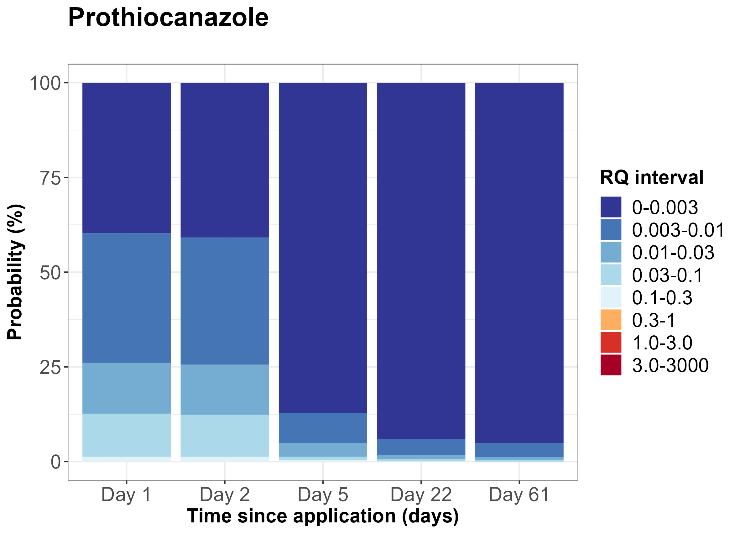

Figure S. 2 Example of clopyralid (a), fluroxypyr-meptyl (b) and prothiocanazole (c) risk estimation over time for 1, 2, 5, 22, and 61 days after application, for the Baseline application scenario, climate model C1 and the time interval of 2070-2100 for NOEC based effect distribution

4.2 **Risk quotient distribution for climate model 1 and application scenarios**

a)
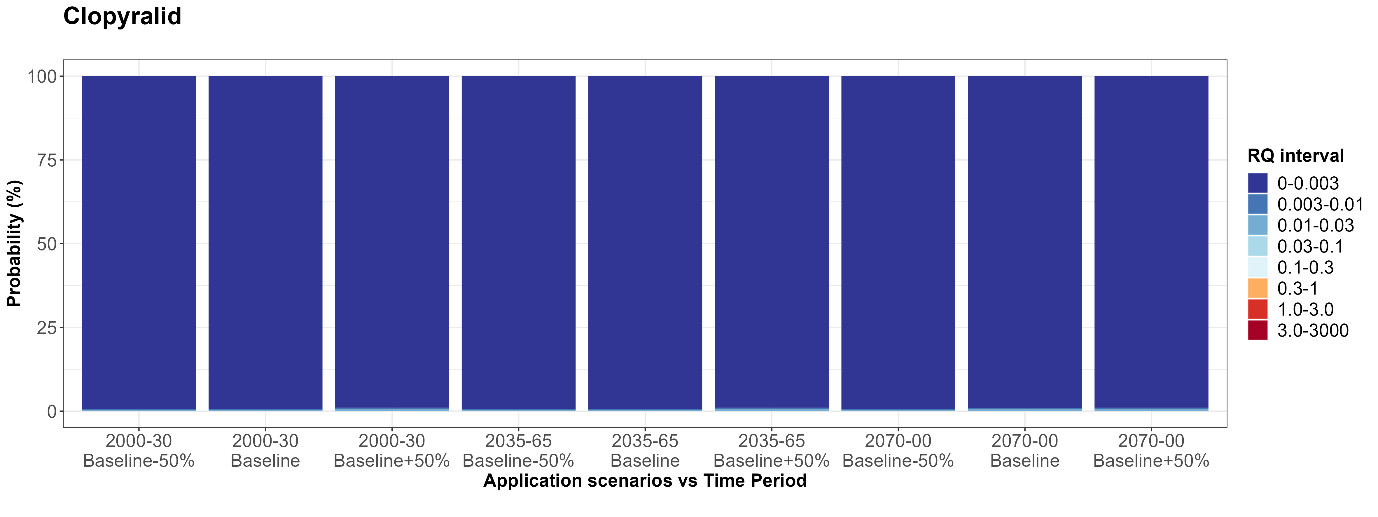

b)
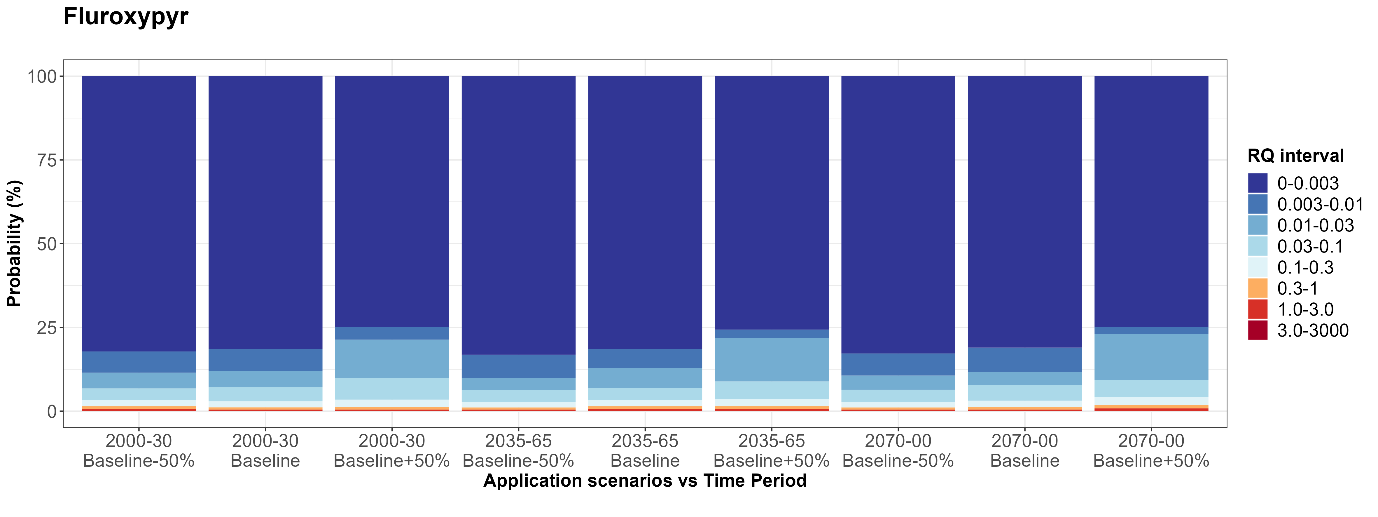

c)
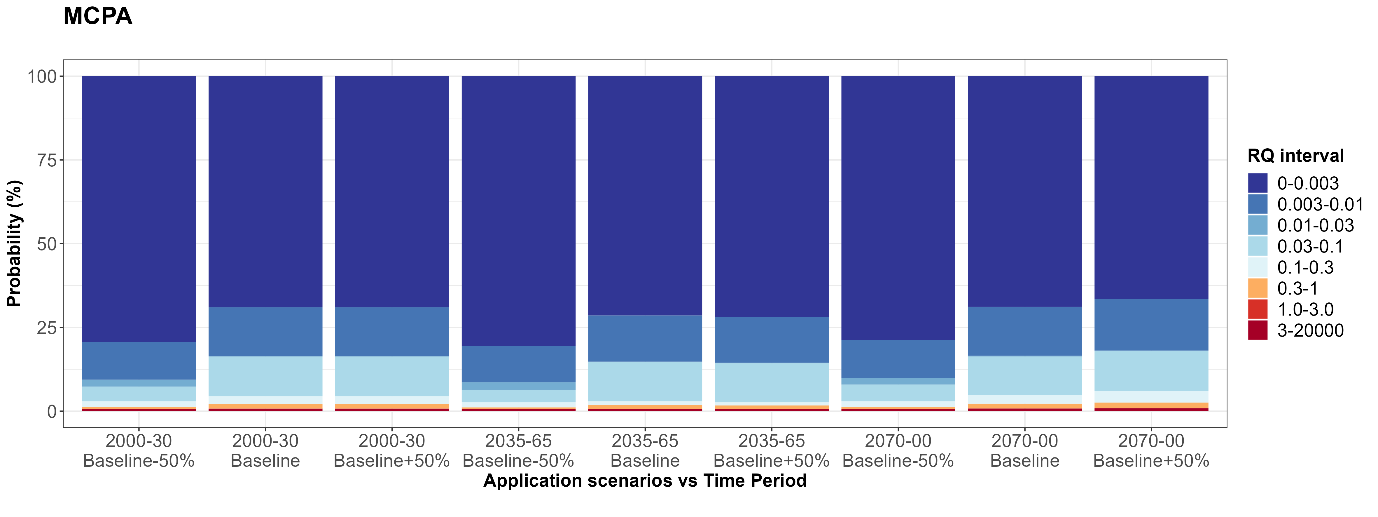

d)
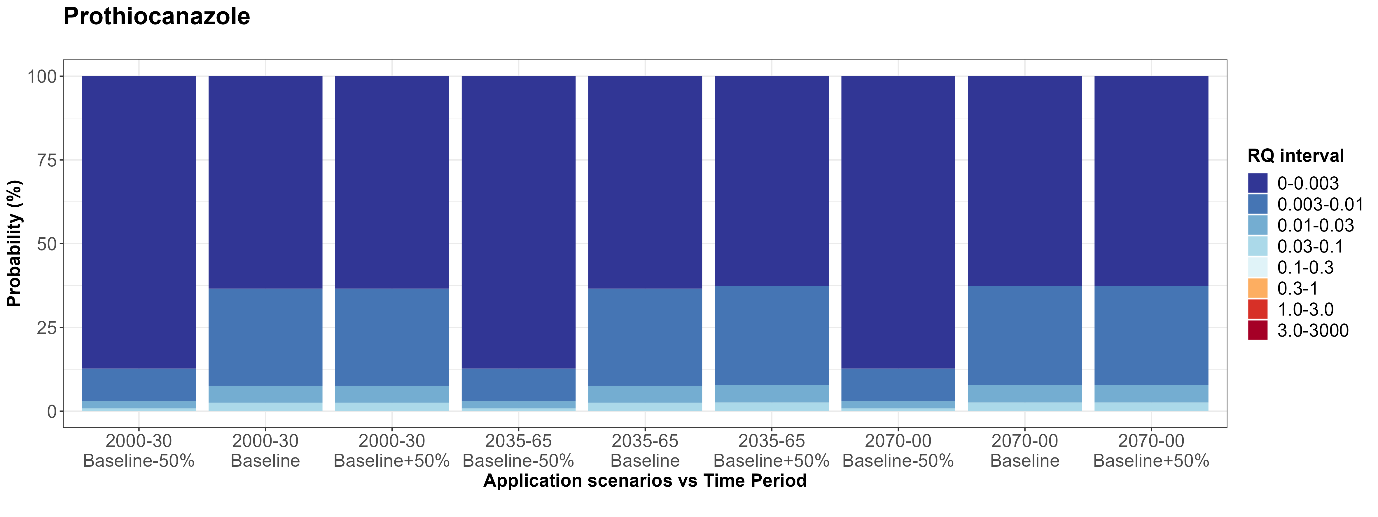
e)
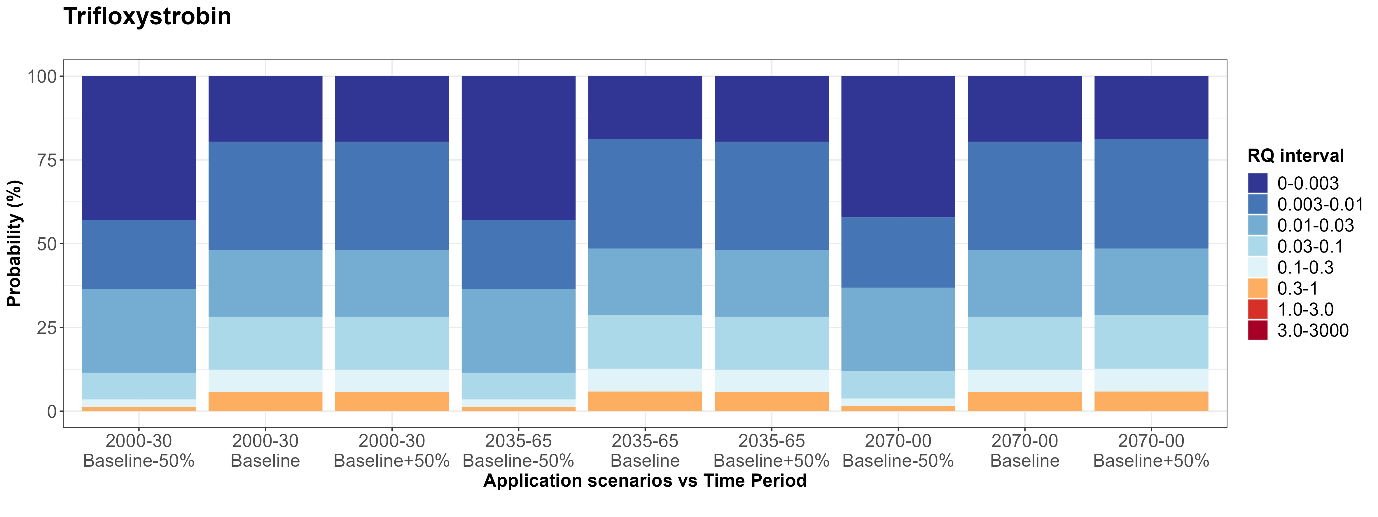

Figure S. 3 Risk estimation of a selected herbicides, clopyralid (a), fluroxypyr-meptyl (b), MCPA (c) and fungicides, prothiocanazole (d) and trifloxystrobin €, for a time since application of 1 day, for the climate model C1 and for EC50 based effects distribution, for all time period and application scenarios.
